# A molecular docking study of human STEAP2 for the discovery of new potential anti-prostate cancer chemotherapeutic candidates

**DOI:** 10.1101/2021.09.02.458478

**Authors:** Timothy Ongaba, Christian Ndekezi, Nana Jaqueline Nakiddu

**Affiliations:** Department of Biomolecular Resources and Biolaboratory Sciences, College of Veterinary Medicine, Animal Resources and Biosecurity (CoVAB), Makerere University, Kampala, Uganda.; Uganda Virus Research Institute,Entebbe, Uganda.; Joint Clinical Research Centre, Kampala Uganda; College of Health Sciences (CHS), Makerere University

## Abstract

Prostate cancer refers to uncontrolled abnormal cell growth (Cancer) within the prostate gland. The disease is a rising health concern and accounts for 3.8% of all cancer deaths globally. Uganda has one of the highest incidence rates of the disease in Africa being 5.2% and many of the diagnosed patients are found to have advanced stage prostate cancer. This study aimed to use STEAP2 protein (prostate cancer specific biomarker) for the discovery of new lead compounds against prostate cancer.

To determine the most likely compound that can bind to STEAP2 protein, we docked the modelled STEAP2 3 Dimension structure against 2466 FDA (Food and Drug Administration) approved drug candidates using Autodock vina. Protein Basic Local Alignment Search Tool (BLASTp) search, Multiple Sequence Alignment (MSA), and phylogenetics were further carried out to analyse the diversity of this marker and determine its conserved domains as suitable target regions.

Six promising drug candidates (ligands) were identified of which Triptorelin had the highest binding energy (-12.1 kcal/mol). Leuprolide was the second most promising candidate with a docking energy of -11.2 kcal/mol. All the top 3 of 6 drugs interacted with highly conserved residues Ser-372 and Gly-369 in close proximity with the iron binding domain that was shown to be important for catalysis of metal reduction.

The two drugs had earlier been approved for treatment of advanced prostate cancer but with an elusive mode of action. Thanks to this study we now have an insight on how their interaction with STEAP2 might be important during treatment.

**Author summary:** Previous studies on prostate cancer prevalence have shown that Uganda among other low to middle income countries has a high prostate cancer incidence rate. Majority of the patients at the time of diagnosis have advanced stage disease. While there has been great improvement in therapeutic options for prostate cancer over the last decade, the drugs currently used are still limited and not 100% effective making cure elusive. This is further compounded by undesirable side-effects from some of the treatments. Altogether this creates a need for the discovery of novel, safe, and efficacious chemotherapeutic agents with minimal or no side effects. In this study, we used a prostate cancer specific protein biomarker; STEAP2 that is highly expressed at all disease stages and is androgen independent and FDA approved drugs from the drugbank to help improve drug specificity, efficacy, and reduce undesirable side effects through *in-silico* methods. The results from the study give an insight on mechanisms of action of current therapy for advanced disease and suggest these very drugs be used even at early stage disease.

## Introduction

Cancer is a disease in which abnormal cells divide uncontrollably and in the process destroy body tissue. It is the second leading cause of death before age 70 years in 91 of 172 countries according to the World Health Organisation in 2015 (1). It is responsible for an estimated 9.6 million deaths in 2018 (1). One in 5 men and 1 in 6 women develop cancer in their lifetime while 1 in 8 men and 1 in 11 women die from the disease (1). As of 2018, the global cancer burden had risen to 18.1 million new cases (1). Worldwide, the total number of people alive with a 5-year cancer diagnosis, called the 5-year prevalence, was estimated to be 43.8 million (2). About 70% to 80% of the avoidable cancer mortality occurs in Low to Middle Income Countries (LMICs) (3). Global cancer incidence is on the rise and it is predicted that by 2050, 70% of the 24 million annual cancer diagnoses will be individuals residing in LMICs (4).

Prostate cancer is the second most frequently diagnosed cancer in men globally and accounts for 3.8% of all deaths caused by cancer in men as of 2018 (1,5). In Uganda for a 16-year period from 1991 to 2006 there was an increase in cancer risk particularly for breast and prostate cancer (4.5% annually) (6). The incidence of prostate cancer in Uganda is among the highest recorded in Africa at 39.6 per 100,000 after age-standardizing (6). A recent study on prostate cancer burden puts its annual incidence at 5.2% (7).

Prostate cancer occurs within the prostate. It can be classified into Localized and metastatic disease depending on absence or presence of spread respectively. Localized prostate cancer is classified as high risk based on clinical staging, prostate specific antigen (PSA) levels, and/or the Gleason score (8). Patients with metastatic prostate cancer are at higher risk of disease and death. The disease can be categorized as castration-sensitive (mCSPC) when surgical removal of the testicles halts cancer advancement due to decreased blood testosterone androgen or castration-resistant (mCRPC) when the cancer continues to progress even in the absence of testosterone androgen (8). Majority of mCSPC patients have a high risk of progressing to mCRPC when initial hormonal treatment fails due to resistance (8).

Prostate cancer can be treated in several ways. However, overall disease management depends on stage/severity and type of cancer. Surgery, nanotechnology for controlled drug delivery, monoclonal antibody therapy, hormonal therapy, radiotherapy, and chemotherapy are some of the treatment methods (9,10). Chemotherapy (especially when hormonal treatment fails), surgery and radiotherapy are the most commonly used modalities of therapy.

Six Transmembrane Antigen of the Prostate 2 (STEAP2) also known as the Six Transmembrane Protein of prostate 1 (STAMP1) is a member of the metalloreductase family that are important in metal reduction of copper and iron. *In vitro* and *in vivo* studies show that STEAP2 plays a key role in prostate cell ability to advance from localised cancer to metastasis (11). STEAP2 is located on the plasma membrane of prostate cells and Golgi complex. It increases prostate cancer progression, controls cell proliferation, differentiation and decreases apoptosis (12). Immunohistochemical staining significantly demonstrate its expression at the cell-cell junctions of prostate cancer cells (13). It is differentially expressed in normal and cancerous tissue making it a potential target for new therapeutic strategies for disease treatment (14). STEAP2 is expressed more than 10 times in normal prostate than other tissues and is exponentially expressed in malignant prostate cancer cells (15,16). It is expressed in low amounts in other tissues such as the brain, liver, skeletal muscle, mammary glands, kidney, lung, uterus, and trachea (15,16). Its levels in these tissues is so low compared to that in the prostate that it is unlikely to have any functional significance (16).

This study aimed at using the STEAP2 prostate cancer biomarker as a target for improving current disease treatment regimen using *in-silico* methods and promising results were achieved.

## Results

### STEAP2 relative tissue expression

The bar graphs figs.1 to 4 show the differential expression of STEAP2 in different tissues as identified by different tissue gene expression software from the Human Protein Atlas. (https://www.proteinatlas.org/). Messenger Ribonucleic Acid (mRNA) expression analysis of the FANTOM5 dataset showed that STEAP2 was most abundantly expressed in prostate tissue followed by ovarian and vaginal tissue (Fig. 2) (17,18). Fig. 2 showed that prostate tissue had the most STEAP2 in mean protein coding transcripts per million as compared to all other listed body tissues using the HPA dataset (18,19).

**Fig. 1:**
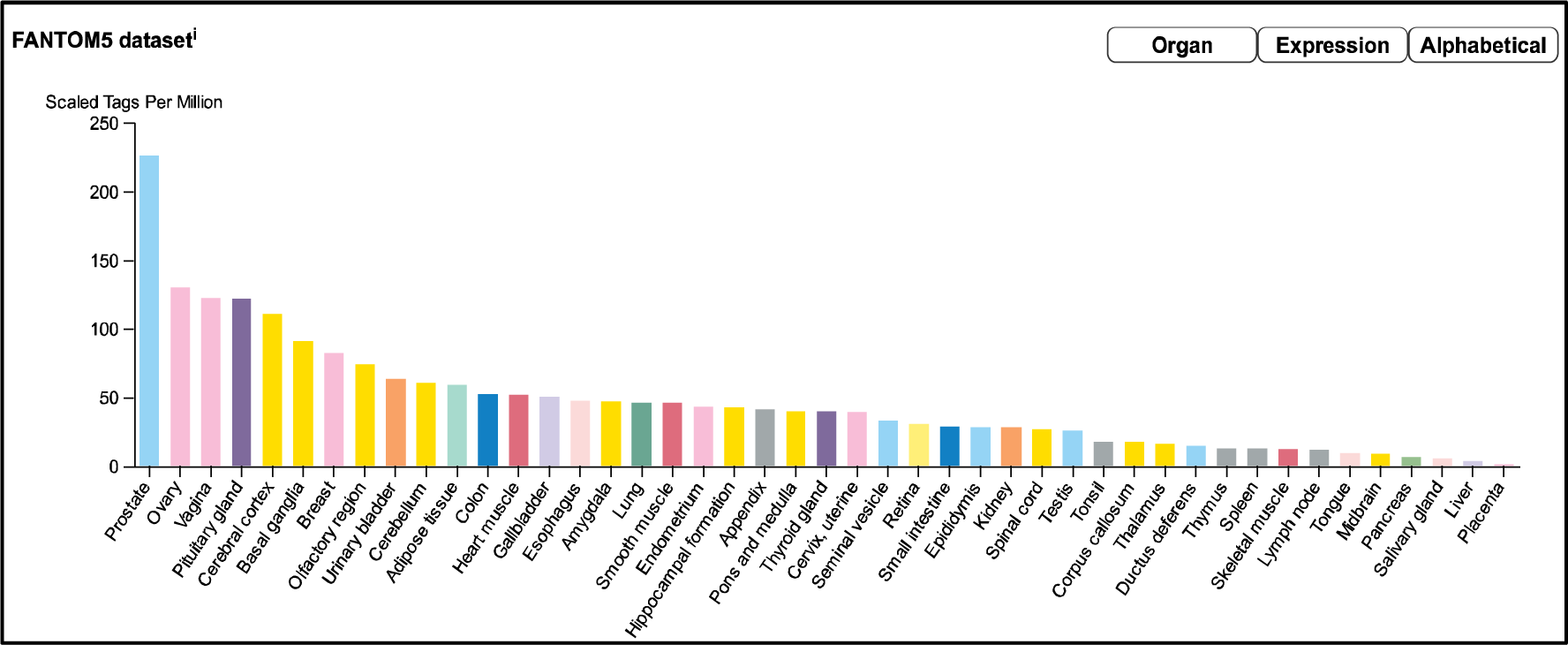
A bar graph showing the differential tissue expression of STEAP2 RNA tissue data reported as scaled tags per million by FANTOM5 dataset (source: https://www.proteinatlas.org/ENG00000157214-STEAP2). The graph shows that STEAP2 mRNA is most abundantly expressed in prostate tissue followed by ovarian and vaginal tissue

**Fig. 2:**
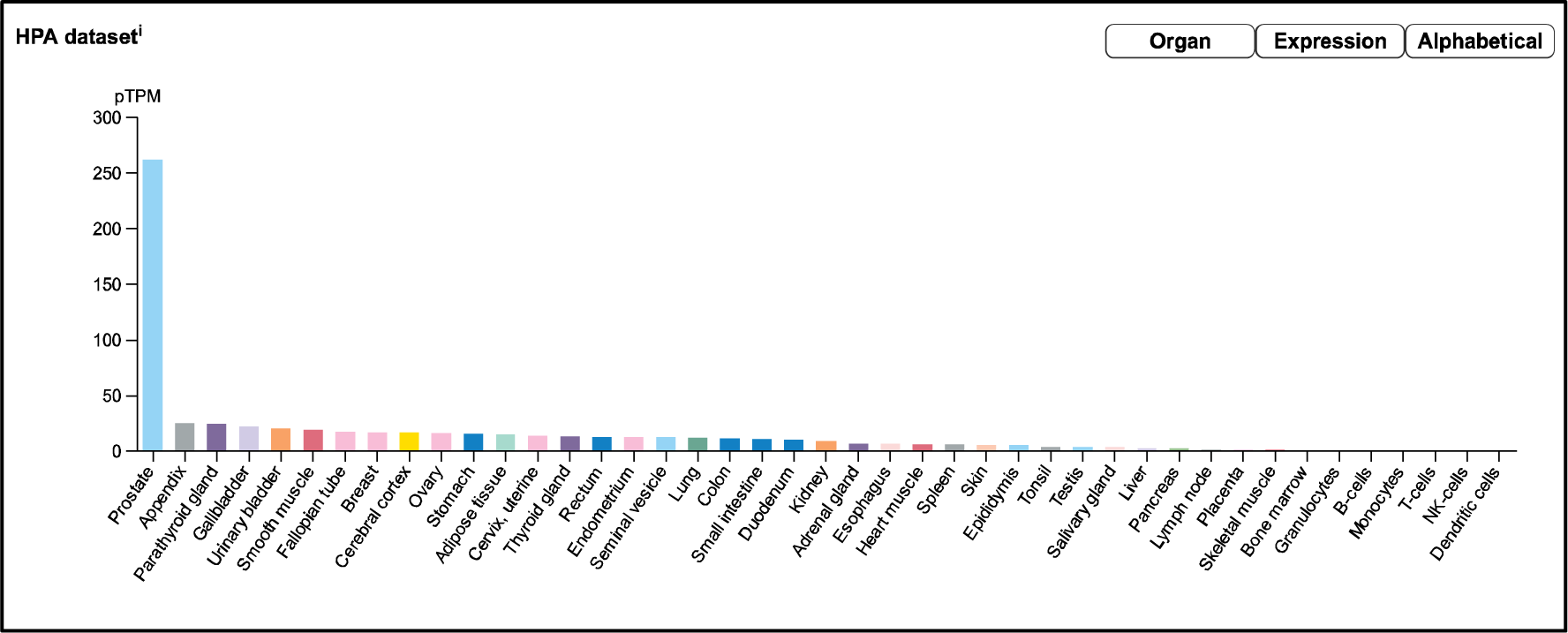
A bar graph showing the differential tissue expression of STEAP2 in mean protein coding transcripts per million by HPA dataset (source: https://www.proteinatlas.org/ENG00000157214-STEAP2). The prostate tissue had a significantly high STEAP2 in mean protein coding transcripts per million as compared to all other listed body tissues followed by the appendix and parathyroid gland.

Fig. 3 further showed prostate tissue to have the highest quantity of STEAP2 mRNA transcripts from different tissue in mean protein coding transcripts per million in comparison to 33 other listed tissues using the GTex dataset (18,20). Fig. 4 combined data from the three datasets by a normalised eXpression level and showed the same trend.(17–20).

**Fig. 3:**
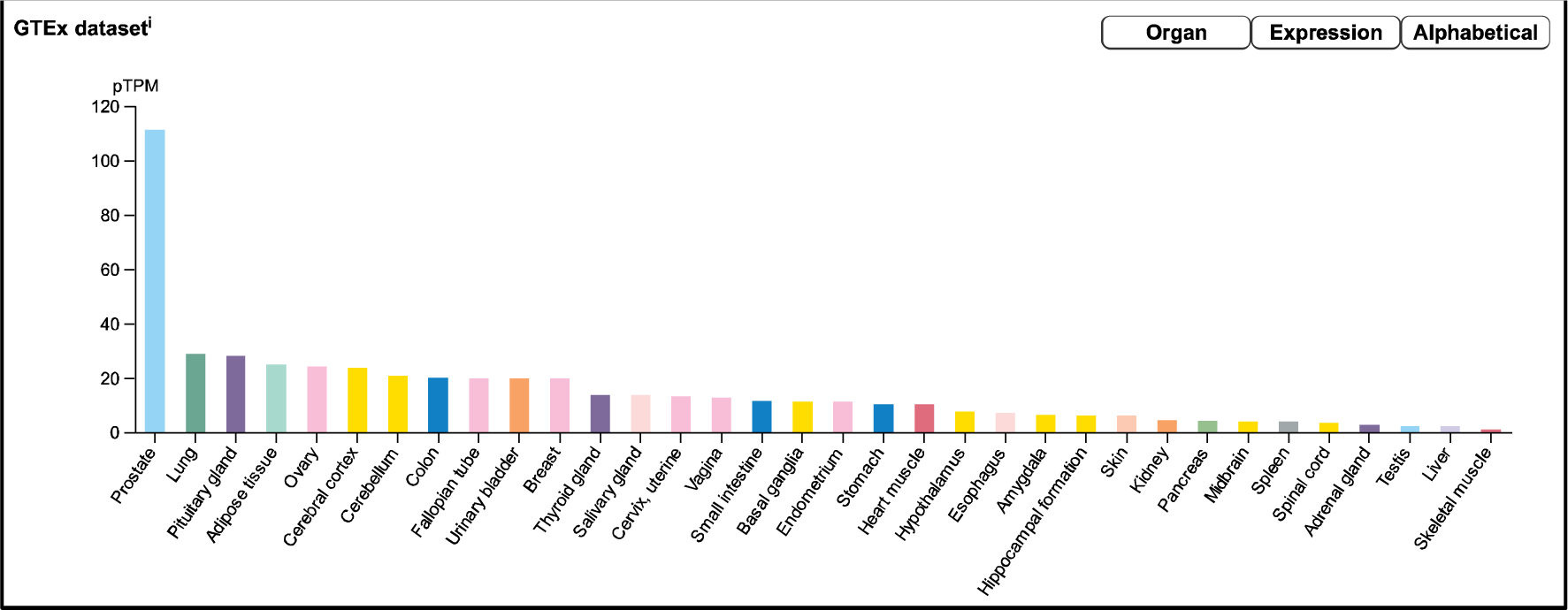
A bar graph showing the differential tissue expression of STEAP2 by RNA sequence tissue reported in mean protein coding transcripts per million shown by GTex dataset (source: https://www.proteinatlas.org/ENSG00000157214-STEAP2/tissue). Prostate tissue has a highly significant content of STEAP2 RNA compared to all other tissues followed by lung and pituitary glands.

**Fig. 4:**
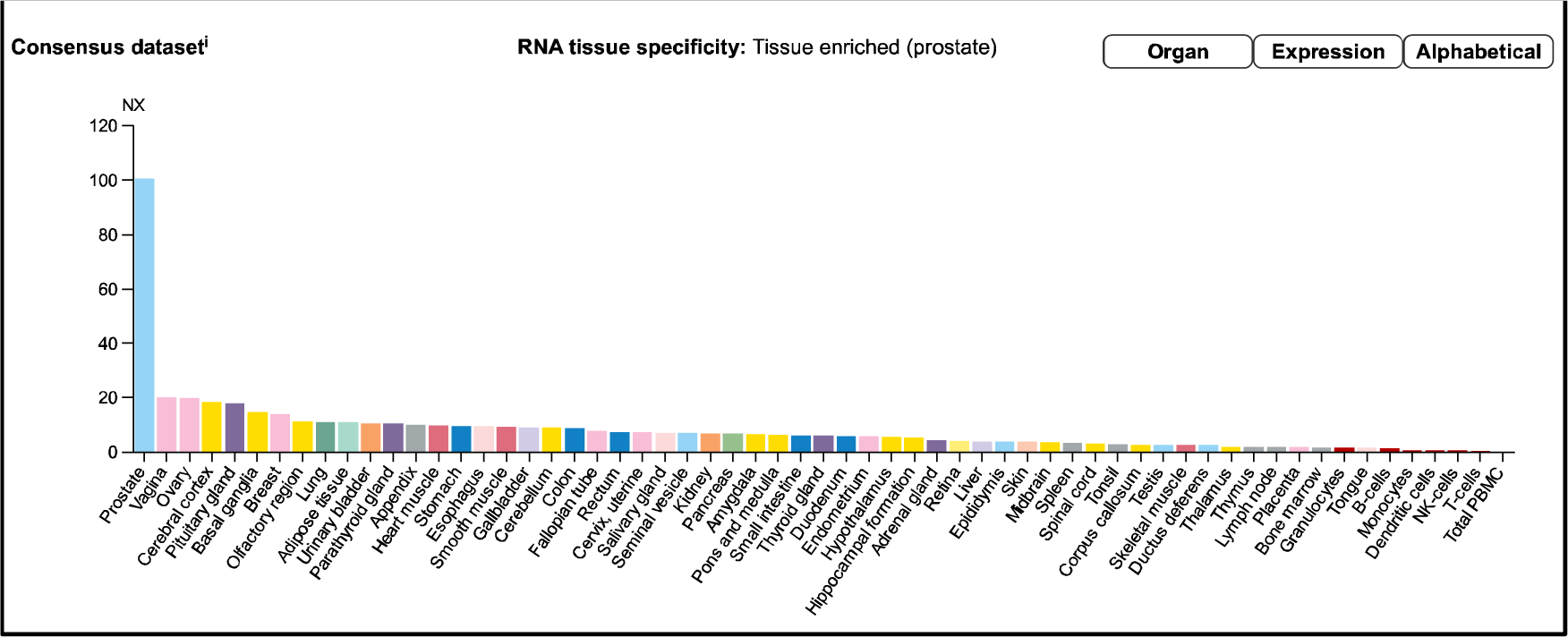
Consensus Normalized eXpression (NX) levels for 55 tissue types and 6 blood cell types, created by combining the data from the three transcriptomics datasets (HPA, GTEx and FANTOM5) using the internal normalization pipeline. (Source: https://www.proteinatlas.org/ENSG00000157214-STEAP2/tissue). The graph maintains the initial observation that prostate tissue contains significantly high amounts of STEAP2 that any other body tissue of the 55 considered.

### Data retrieval of STEAP2 sequence

The protein *H. sapiens* STEAP2’s (Uniprot accession number: Q8NFT2) in FAST All (FASTA) format from UniProt (Bateman *et al.,* 2017);

### Multiple Sequence alignment

A dendogram generated using Molecular Evolution Genetic Analysis (MEGA) showed that STEAP2 was conserved mostly in species under phylum Chordata(Fig. 5) (21) The tree was a result of 1000 bootstrap iterations with the values at each node indicating the percentage times of 1000 that particular node was redrawn as a result of genetic relationship of its taxons.

**Fig. 5.**
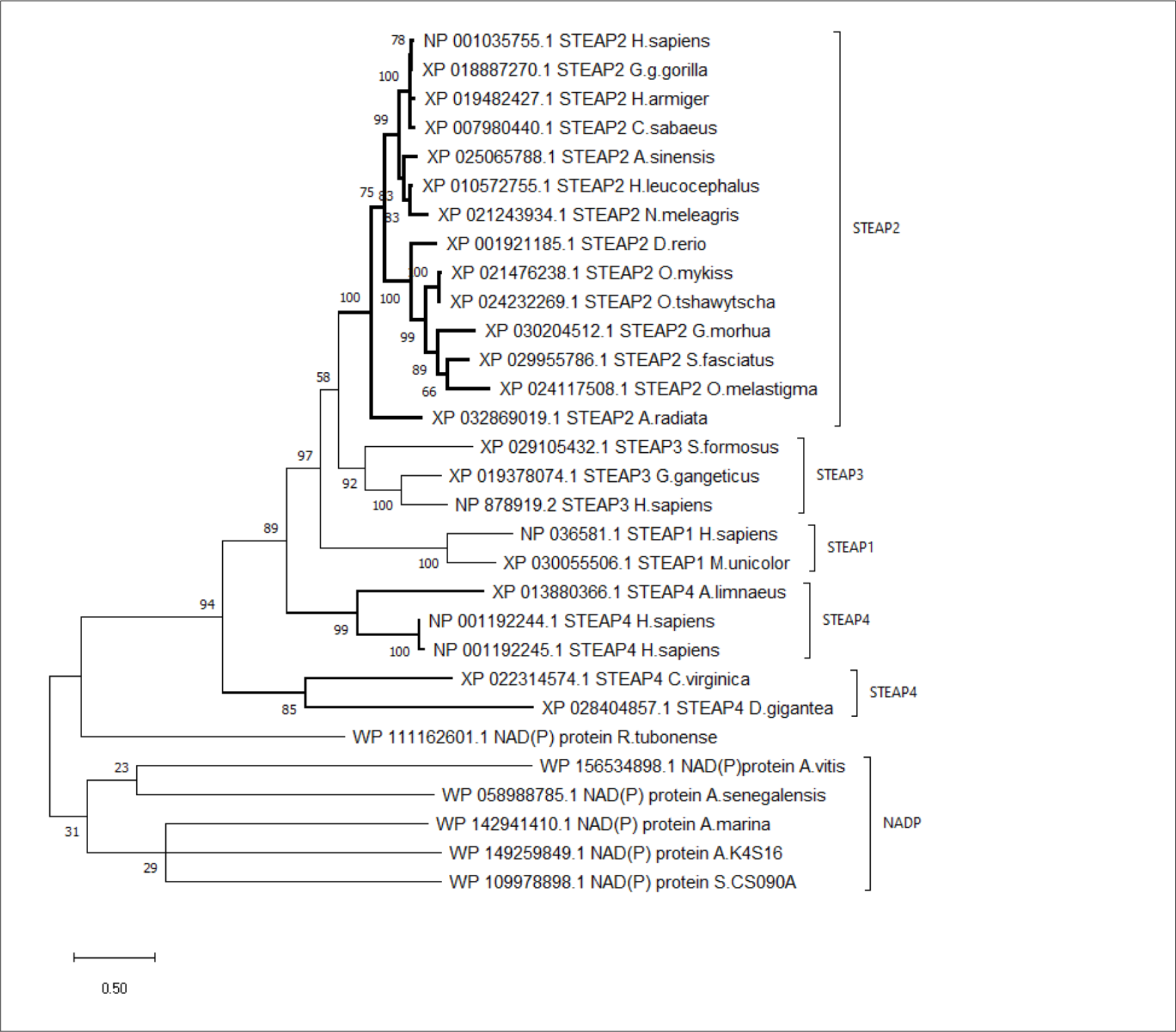
The Phylogenetic tree of H.sapiens STEAP2 shows that it is most related to G.gorilla gorilla since it is redrawn 78% of 1000 times by the bootstrap method. This therefore, indicates that these three species’ have the most conserved regions (21,22)

The phylogenetic tree showed the evolutionary relationship between the 30 selected species in relationship to STEAP2. The highlighted clade shows species containing STEAP2 that also happen to be under phylum Chordata. The species go on differentiating into sub taxa till the mammalian class and primates order where our query species; *H. sapiens* falls together with its most similar species; *G.g.gorilla.* as shown in Fig. 5. The rest of the clades contain other STEAP proteins and one NADP from other phyla.

Having clustered the STEAP2s alone, a second phylogenetic analysis of these was carried out to have a more accurate and precise evolutionary relationship between the different species’ proteins (Fig. 6).

**Fig. 6:**
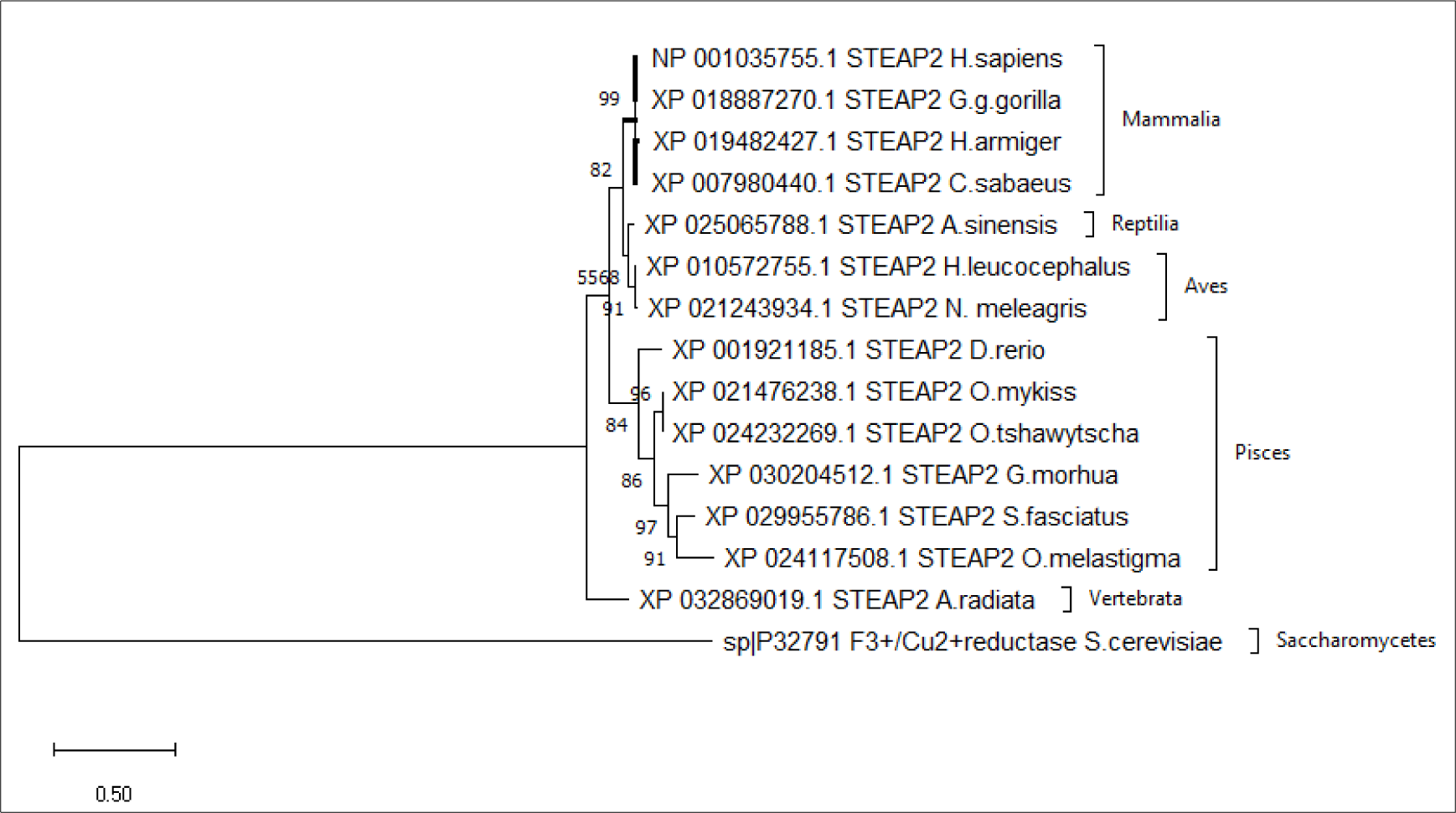
From the phylogenetic tree, which is a result of 1000 bootstraps, it is shown that the STEAP2 protein from G.g.gorilla (mammal) is most related to human STEAP2 from 99% of the 1000 redrawn trees and the least related is A.radiata. The first four listed species from top belong to class Mammalia, followed by an alligator (Reptilia), birds (Aves) fish (Pices) and a turtle (vertebrata) all under phylum Chordata.

It is noteworthy that the mammalian species baring STEAP2 clustered together, followed by those of reptilia, pisces, and vertebrata species; *A. radiata* with the tree having been rooted to a yeast species; *S. cereviciae*.

### Analysis of highly conserved residues from the MSA that might be responsible for the phylogenetic relationships of the different species

The MSA image depicts the conserved residues across 15 selected sequences of STEAP2 baring species and S. cerevisiae from the original 30 from the protein BLAST search. The multiple sequence alignment (MSA) was done using MUSCLE in Jalview to determine the most conserved residues among different species and these conserved residues were highlighted in columns (23,24) The first MSA image of the 30 sequences is attached in SI_Fig.17. The colouring was done by clustalx in both images and the length of sequences in the wrap was adjusted to first 100 residue window. The sequences that were used for the MSA are detailed in SI_Table1.

Some of the conserved residues as highlighted in colour in the MSA with respective Weblogos to the left were found to be associated to functional motifs such as the Rossman fold that is closely associated with NADP domain (25) and some of the transmembrane domains were associated with iron binding domains.

### Homology modelling

The query sequence of STEAP2 was retrieved from UniProt in FASTA format (UniProt accession number: Q8NFT2). The template search results by the homology modelling engines together with parameters of similarity to STEAP2 are shown (table 1). Template 6HCY/6HDY (human STEAP4) was the most common top hit.

**Table 1:** Top homology modelling templates from selected model engines and respective parameter of the hits.

| ENGINE | SEQUENCE<br>IDENTITY<br>(%) | COVERAGE | ORGANISM | PDB ID | STOICHIOMETRY | RESOLUTION<br>(Å) | STRUCTURAL<br>DETERMINATION |
| --- | --- | --- | --- | --- | --- | --- | --- |
| HHPRED | 46.00 | 18-171 | *H.sapiens* | 6HCY_A | Homo-3-mer | 3.1 | EM |
| SWISS<br>MODEL | 47.61 | 30-466 | *H.sapiens* | 6HCY.1.A | Homo-3-mer | 3.1 | EM |
| PRIMO | 53.00 | 33-215 | *H.sapiens* | 2VNS | Homo-2-mer | 2.0 | X-RAY DIFFRACTION |
| RAPTORX |  | 31-466 | *H.sapiens* | 6HD1C | Homo-3-mer | 3.8 | EM |
| ROBETTA | 47.83 | 1-490 | *H.sapiens* | 6HCYA | Homo-3-mer | 3.1 | EM |
| I-TASSER | 47.90 | 0.890 | *H.sapiens* | 6HCYC | Homo-3-mer | 3.1 | EM |

Since the engines search the PDB database and with different principles, there was a difference in top hits.

The common hit among 5 of 6 engines was human 6HCY/ 6HD1 (STEAP4) with a better coverage than 2VNS identified by PRIMO (Table 1). In all but PRIMO, the top hit was 6HCY with better coverage than 2VNS as a template. This suggests 6HCY as the best template for homology modelling.

The selected templates were used to model human STEAP2 and the homology models are displayed in Fig. 8 in cartoon format and coloured by chain. The homology model from PRIMO was not satisfactory because its best hit template (2VNS) had a very small template coverage of 183 amino acids out of the 490 amino acid query sequence. PRIMO only provides templates that have been resolved by X-ray crystallography as such, it was not possible to select the 6HCY/ 6HD1 (STEAP4) template. X-ray crystallography is also limited by size in terms of structure determination. It is ideal for small size proteins and not large sized proteins like STEAP4 a homo-3-mer (26).

**Fig. 7:**
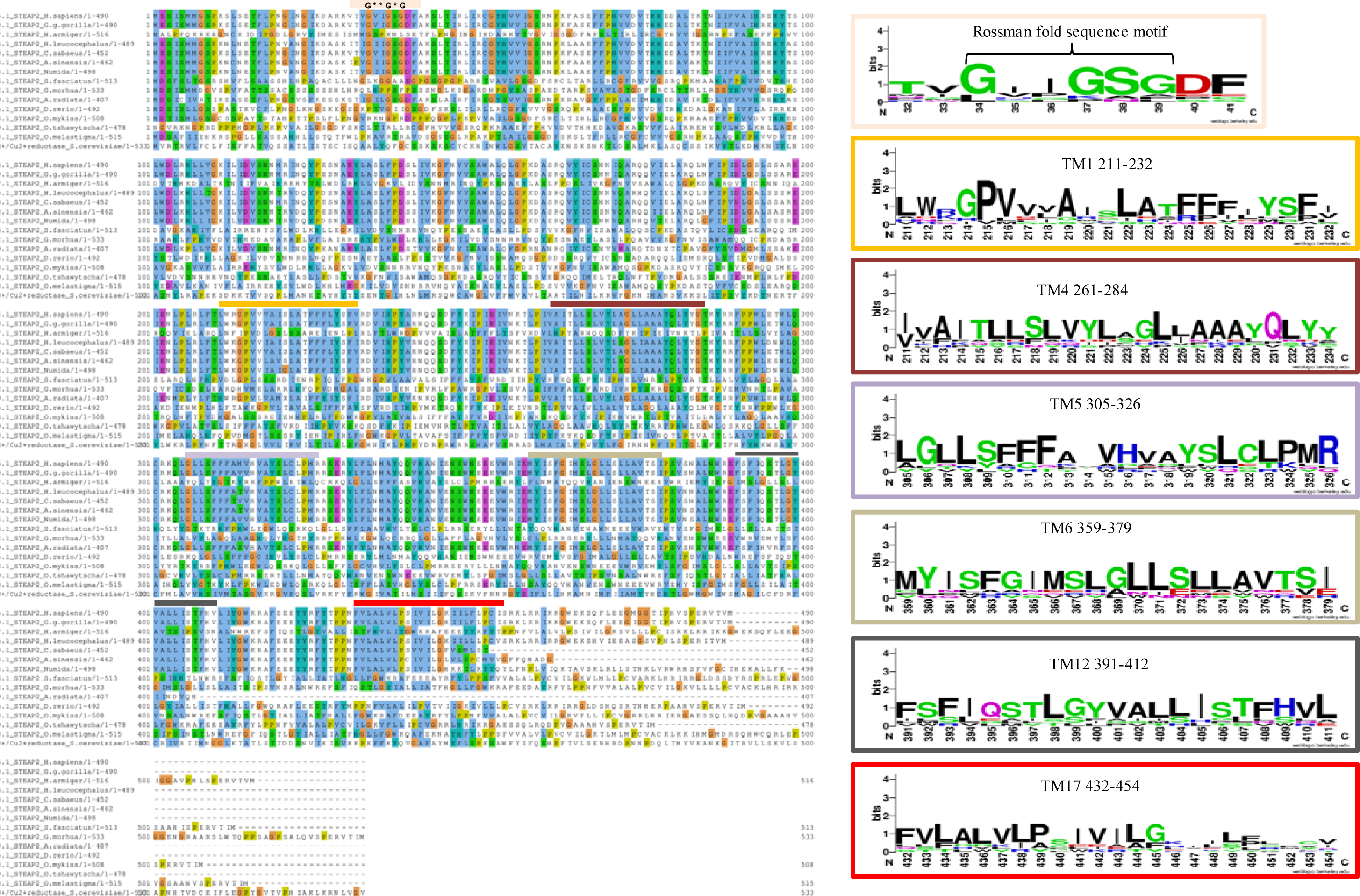
An MSA of STEAP2 proteins from 14 species from 4 different classes of chordates. The most conserved residues are associated with functional roles of the protein. From top, the Rossman fold, an alpha beta alpha secondary structure with sequence GxGxxG is closely associated with the dinucleotide phosphate group of NADP followed by the first trans-membrane domain (TM1) (25). The second, third, fourth and fifth TM domains (TM4,5,6 and 12 respectively) in close contact with the iron binding TM domain and finally the sixth TM domain

**Fig. 8:**
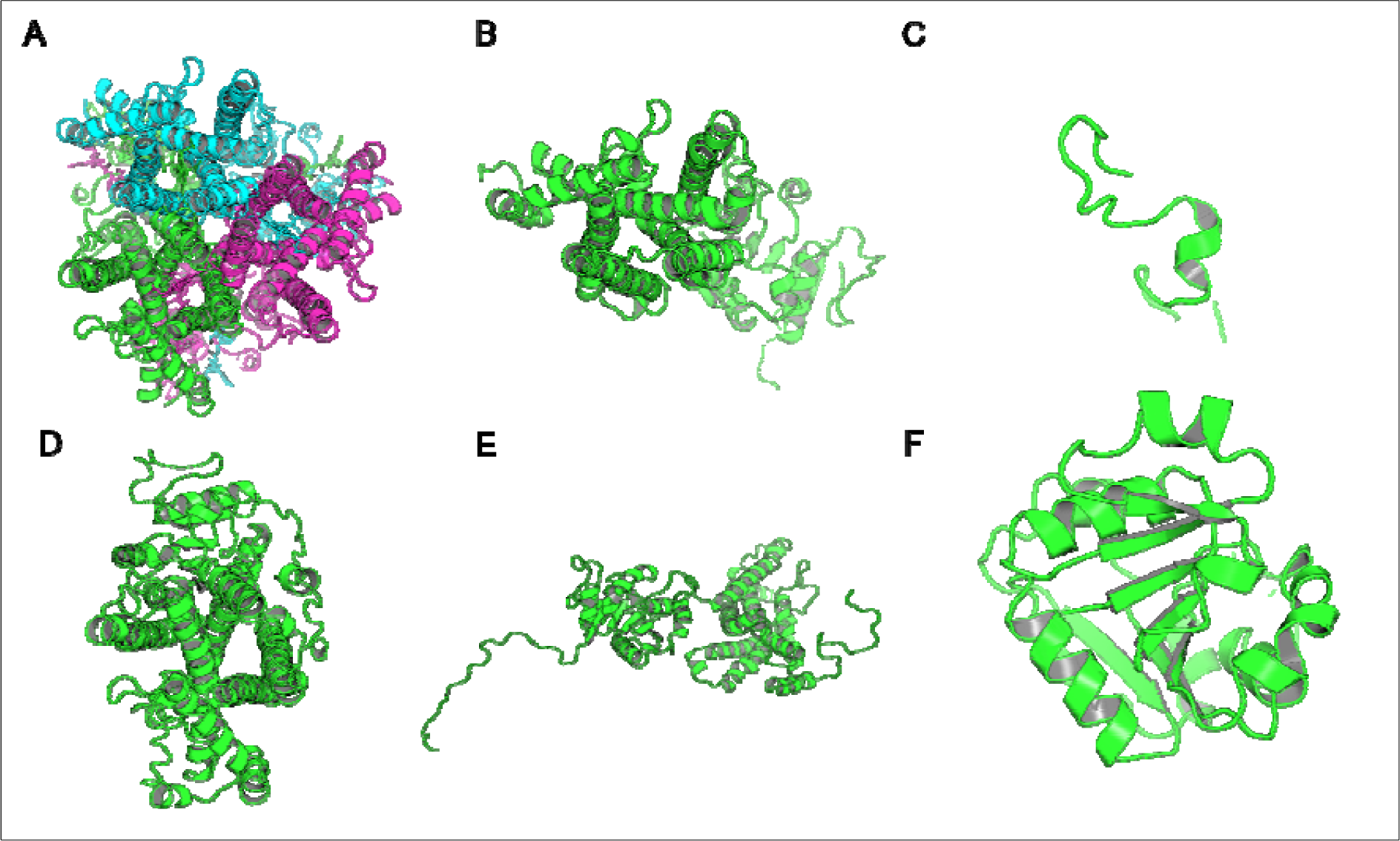
*The images of the homology models were shown in cartoon format and coloured by chain. Image A shows the homology model by SWISSMODEL, B: The top homology model by I-Tasser, C: The homology model by HHPRED, D: The homology model by RaptorX, E: The homology model by Robetta and. F: The homology model by Primo. Only the homology model by SWISSMODEL was a homo trimer, the rest were single chain models and the used template; human STEAP4 is a homotrimer*

Homology models from the different engines (images generated in cartoon format using PyMol).

The homology model from SWISSMODEL (Fig. 8-A) was a homo-3-mer made up of 437 amino acids per chain. The template used was 6HCY and the homology model had the least RMSD of 0.086 when superimposed with the 6HCY template (Table 2). Additionally, on average it had the best score by the used evaluation tools. For example, the modeller normalised DOPE score of -5.95, the Ramachandran plot having 94.4 of its residues lying within the favoured region a -3.06 and-6.33 Z-score from QMEAN and ProSA respectively. Altogether, these results show that the SWISSMODEL homology model was most similar model to the used template (27).

**Table 2:**
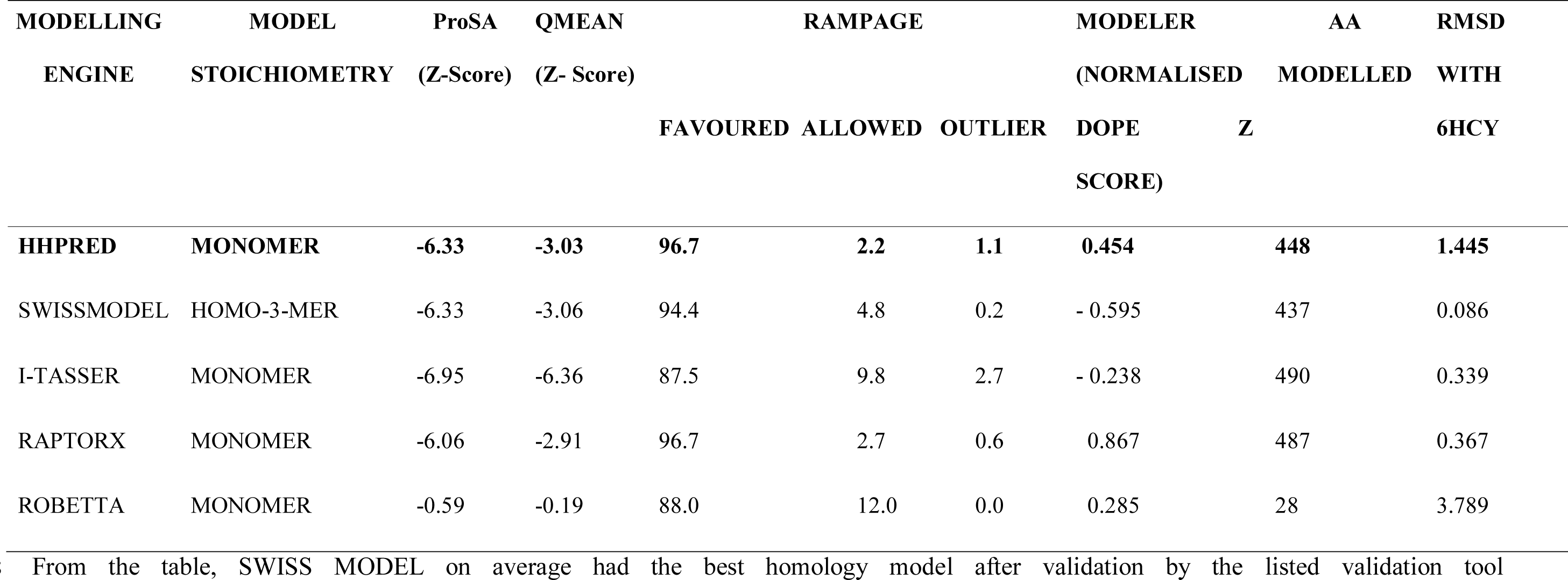
Structural evaluation of homology models by different validation tools.

The I-TASSER model (Fig.8-B) was a monomer of 490 amino acids had a RMSD of 0.339, the second best after that by SWISSMODEL. It had the best ProSA Z score of -6.95 QMEAN Z score of -6.36 but the worst percentage of residues (87.5%) in the favoured region on a Ramachandran plot when compared to other homology models. It also had the second best Modeller normalised DOPE score of -.0238. It had no long ending loops and was therefore of satisfactory quality.

The HHPRED model (Fig.8-C) was a monomer of 448 amino acids. The used template was 6HCY which on superimposition with the homology model had a 4^th^ best of 5 RMSD of 1.445. It had a second best ProSA Z score of -6.33 same as that for SWISSMODEL, a third best -3.03 QMEAN Z score, the best percentage of residues falling in the favoured region on a Ramachandran plot (96.7) but had the second last Modeller normalised DOPE score of 0.454. It had long loops which are indicators of non-satisfactory regional model quality.

RAPTORX model (Fig.8-D) was a monomer of 487 amino acids, had a third best RMSD of 0.367. It had the best percentage of residues within the favourable region on a Ramachandran plot together with that from HHPRED. It however, had the second last of ProSA and QMEAN Z scores, -6.06 and -2.91 respectively and the last Modeller normalised DOPE score of 0.867. It also had noticeable long loops at the extremes of its upper and lower domains.

ROBETTA model (Fig.8-E) was a monomer of 28 amino acids. It was too short, a product for a sequence of 490 residues. It had a 3^rd^ best Modeller normalised DOPE score of 0.285, a 4^th^ best 88% percentage of residues falling within the favoured region on a Ramachandran plot. It however, had the last of 5 for the RMSD, ProSA and QMEAN Z scores that were 3.789, -0.59 and -0.19 respectively. It was, therefore, not a satisfactory homology model.

The model by PRIMO (Fig.8-F) was a monomer of 183 amino acids. This was short a homology model for a query sequence of 490 amino acids. It was not considered for further evaluation as the other 5 because its template was already different from them.

The above stated results are summarised in table 2.

Fig.9 showed the ProSA evaluation of SWISSMODEL which was the best homology model together with its Ramachandran plot on the right. The z-score of -6.33 refers to the standard deviation of the homology model from an ensemble of misfolded proteins whose structures were determined by NMR and X-ray crystallography (28). The Ramachandran plot shows a plot of the homology model’s residue in which those in black, dark grey and light grey regions represent highly preferred conformations (Favored), white with black grid region represents prefered conformations (Allowed) and white with grey grid lines region represents questionable conformations (outlier).

**Fig.9.**
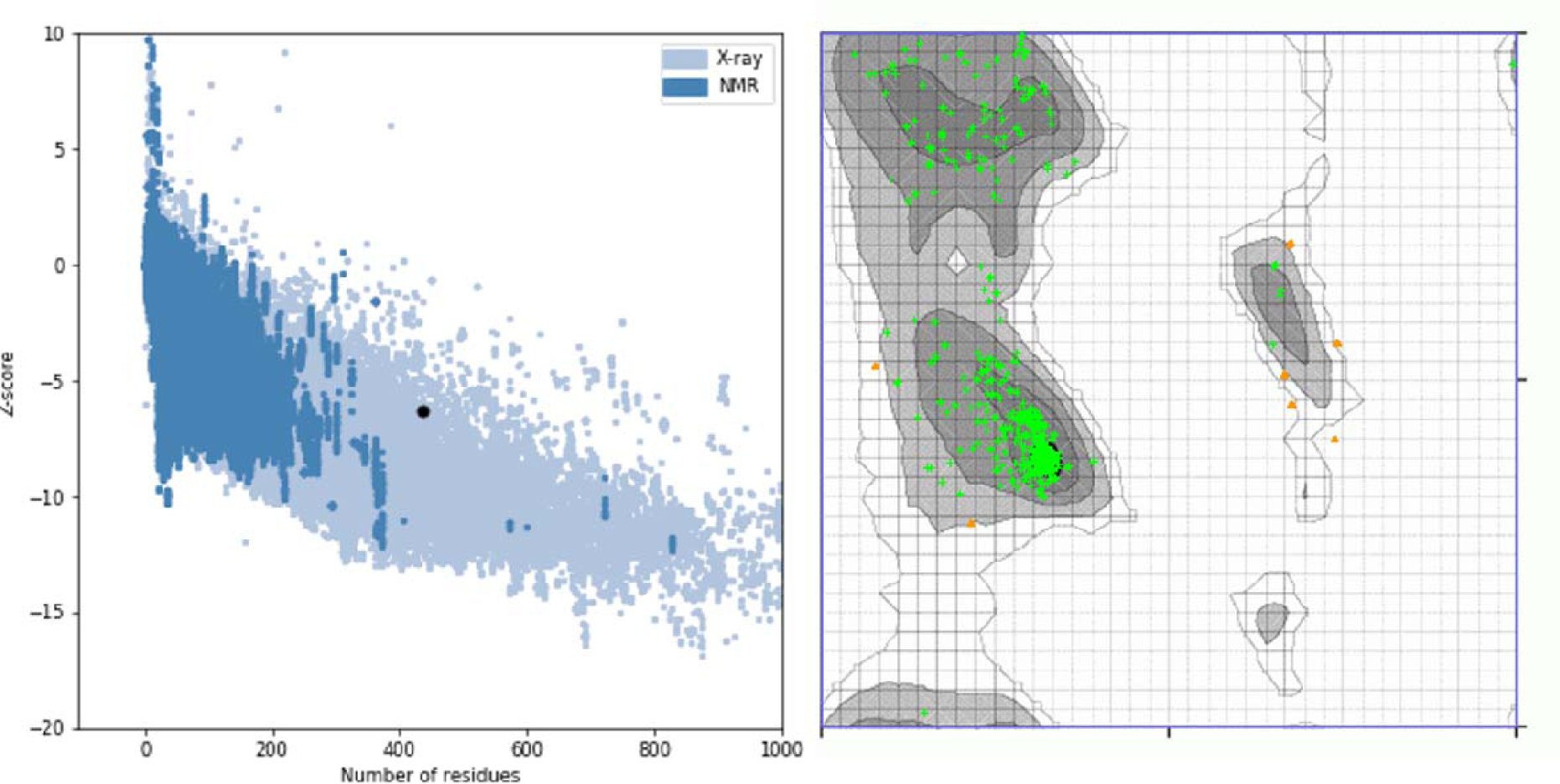
A ProSA z-score and Ramachandran plot of the SWISSMODEL homology model. The z-score of -6.33 suggests that the homology model is properly folded and 96.7% of the protein residues being under the favoured region shows that almost all the protein residues are properly sterically placed in 3D space.

**Fig.10:**
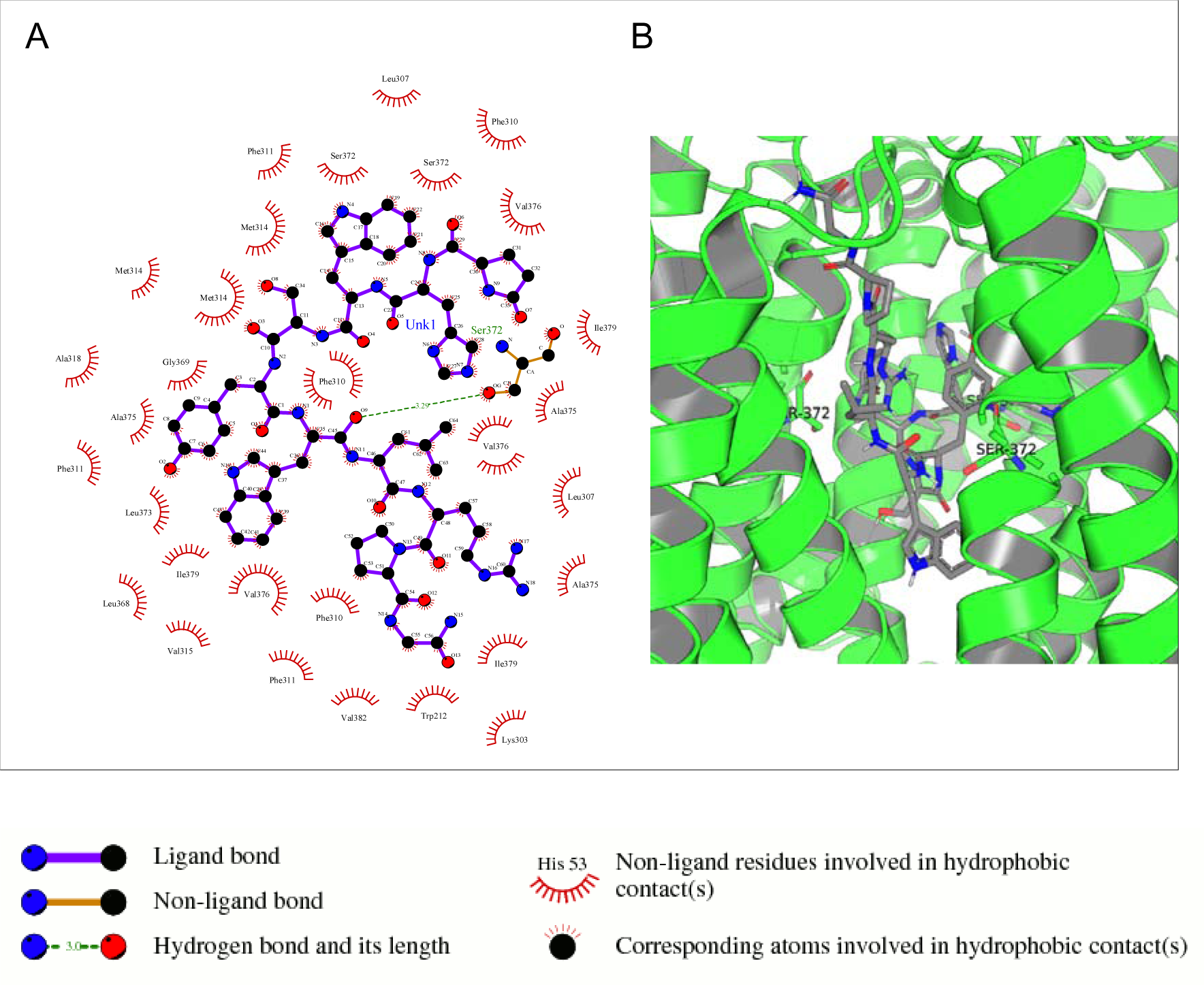
A protein-ligand complex of STEAP2 and Drugbank1543 shows residue Ser-372 from the 6^th^ TM domain according to the MSA of STEAP2s and this forms a hydrogen bond with an atom of the ligand (Drugbank1543). The rest of the interacting receptor residues are shown as crescents and these form other bonds with the receptors. These interacting residues are many compared to other complexes and contribute a significant binding energy of the complex.

### Molecular Docking

Figs 11 to 16 show the interaction of six different residues and part of the receptor molecule STEAP2. The types of bonds between interacting residue and ligand atoms such as hydrogen bonds and other forces of attraction between different atoms are shown in the 2 dimension ligplots as shown by the key. To the right, the 3-dimension image shows the ligand with the respective interacting receptor residues that form hydrogen bonds with the respective ligands labelled and shown as sticks.

**Fig.11:**
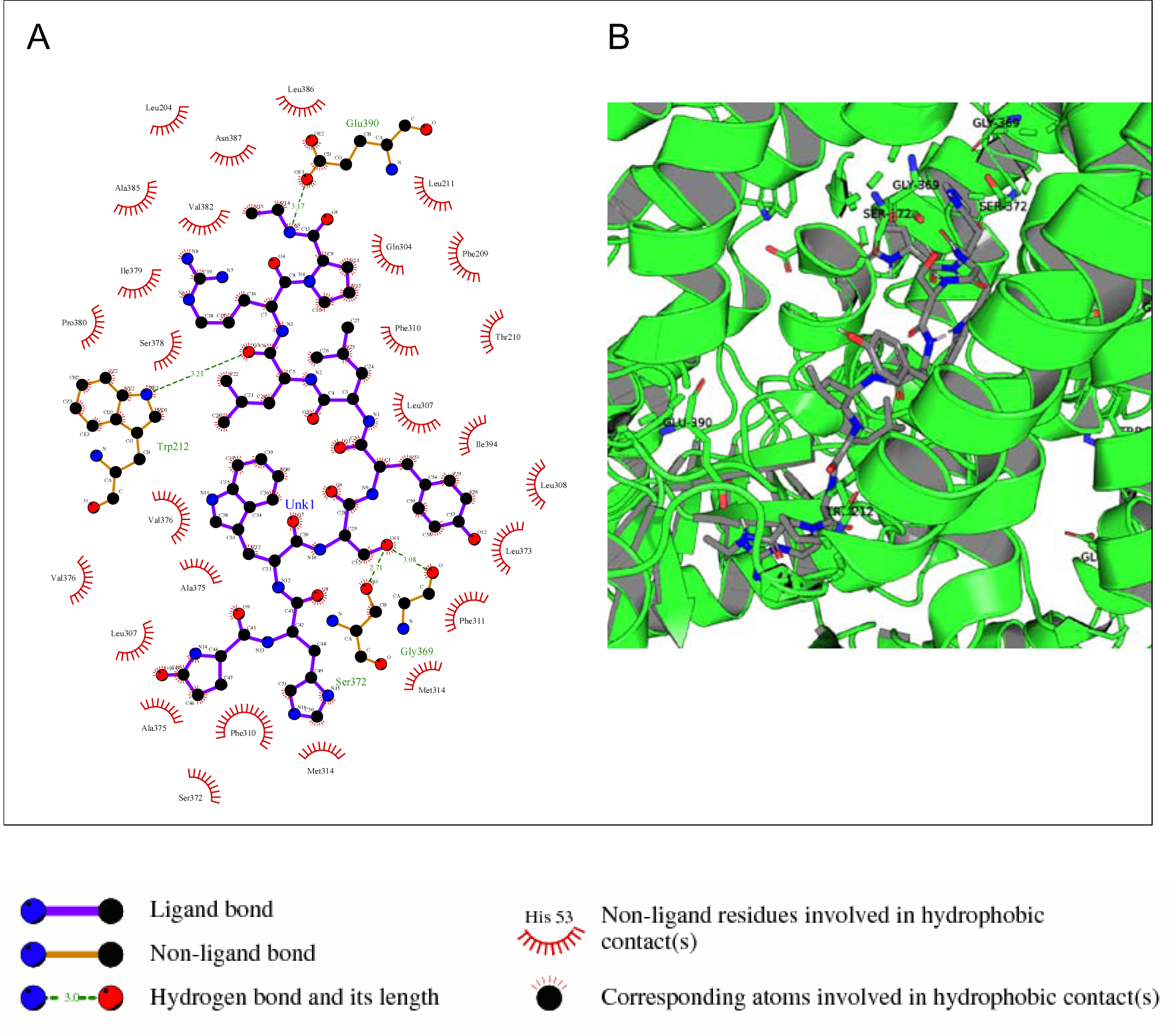
A protein-ligand complex of STEAP2 and Drugbank2 shows residues Glu-390, Gly-369, Ser-372 and Gly-369 within the 6^th^ and 12^th^ TM domain form hydrogen bonds with ligand Drugbank2.

**Fig.12:**
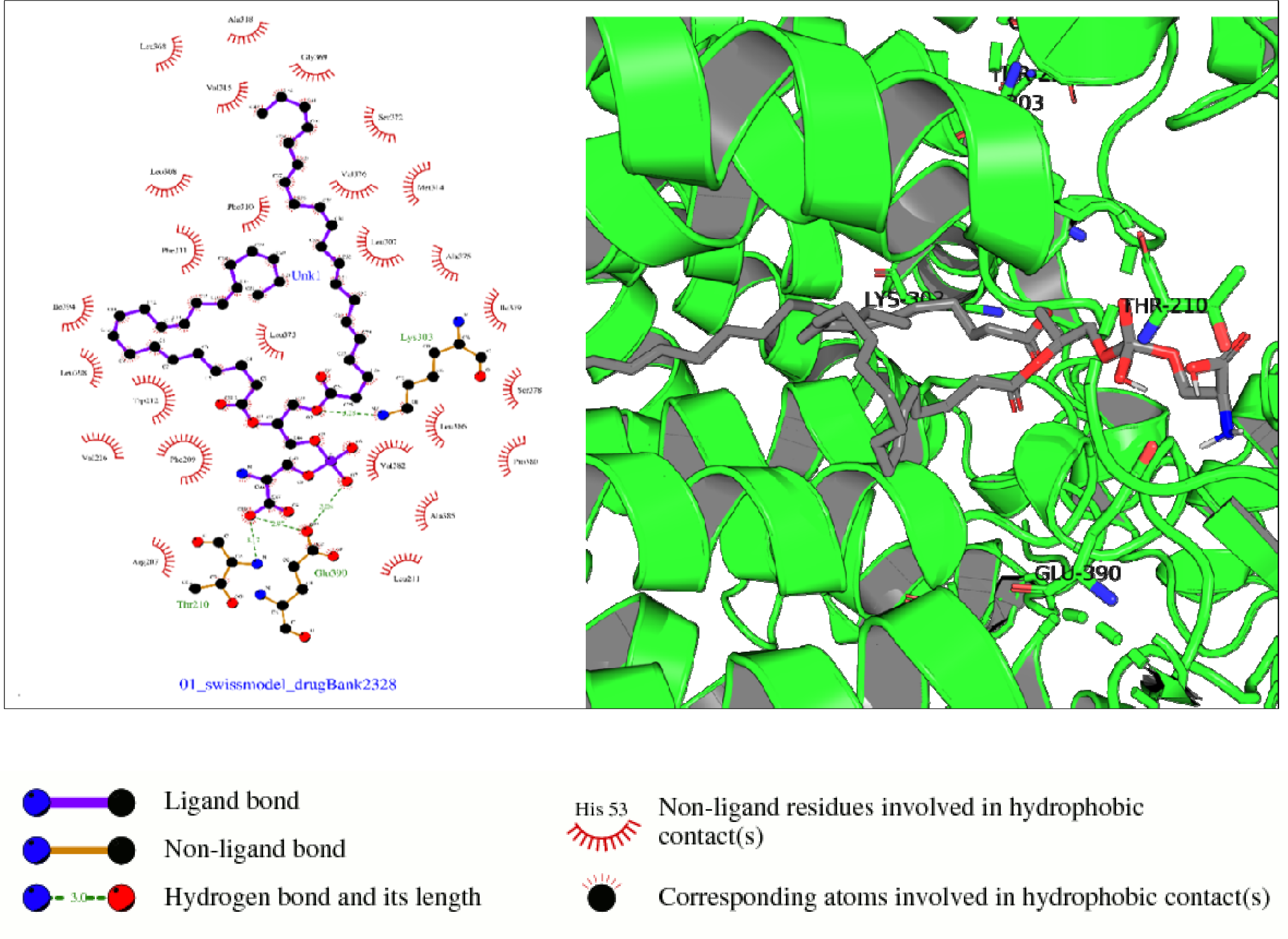
In the above complex, receptor residues Thr-210 (NADP domain), Glu-390, and Lys-303 (TM5) form hydrogen bonds with ligand Drugbank2328.

**Fig.13:**
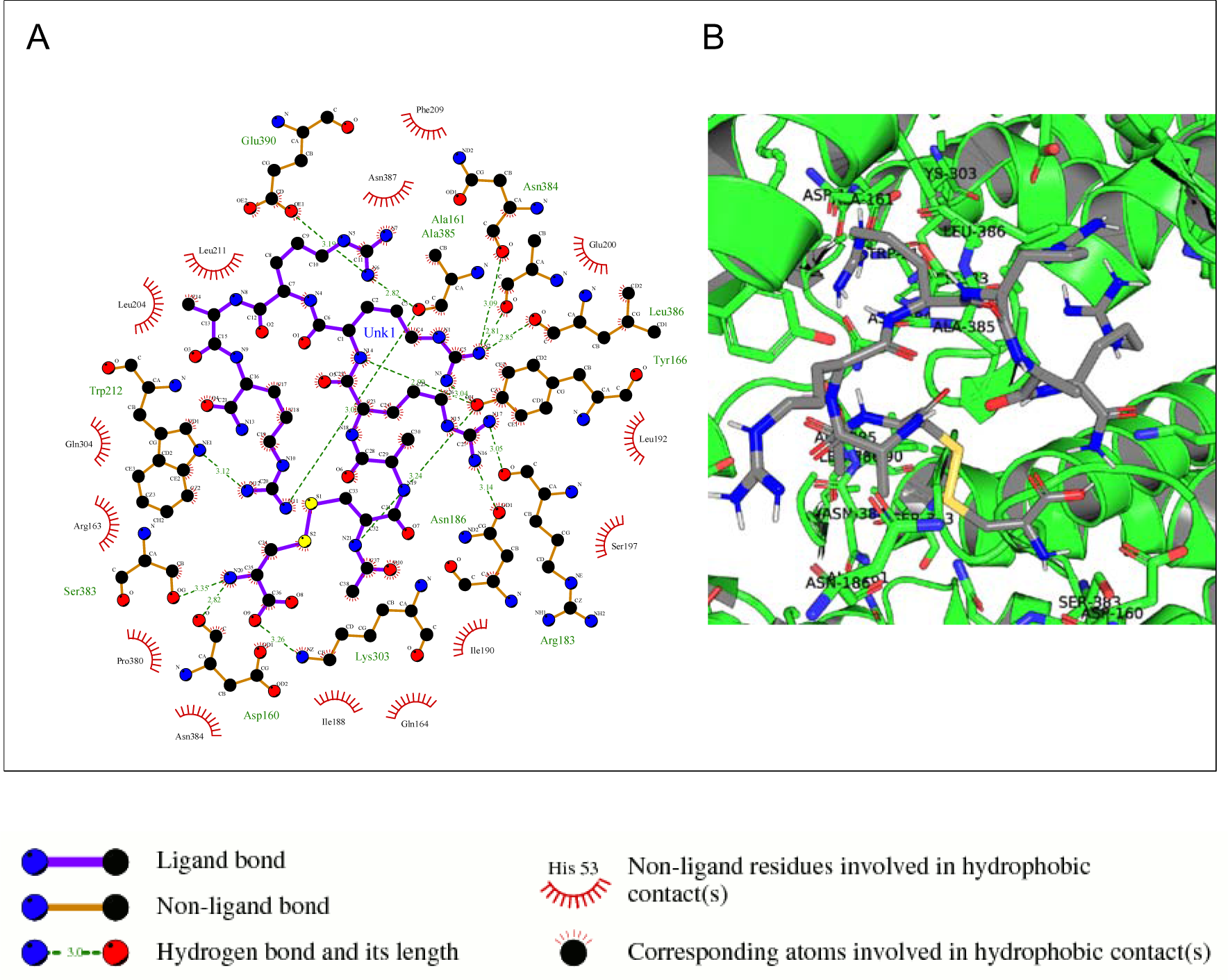
In *the above complex, receptor residues Glu-390, Ala-161,386, Asn-186,384, Lue-386, Tyr-166, Trp-212, Ser-383, Lys-303, Arg-183 and Asp160 all form hydrgen bonds with ligand DrugBank2154. The residues are majorly from the NADP binding dominan and the 12^th^ TM domain*

**Fig.14:**
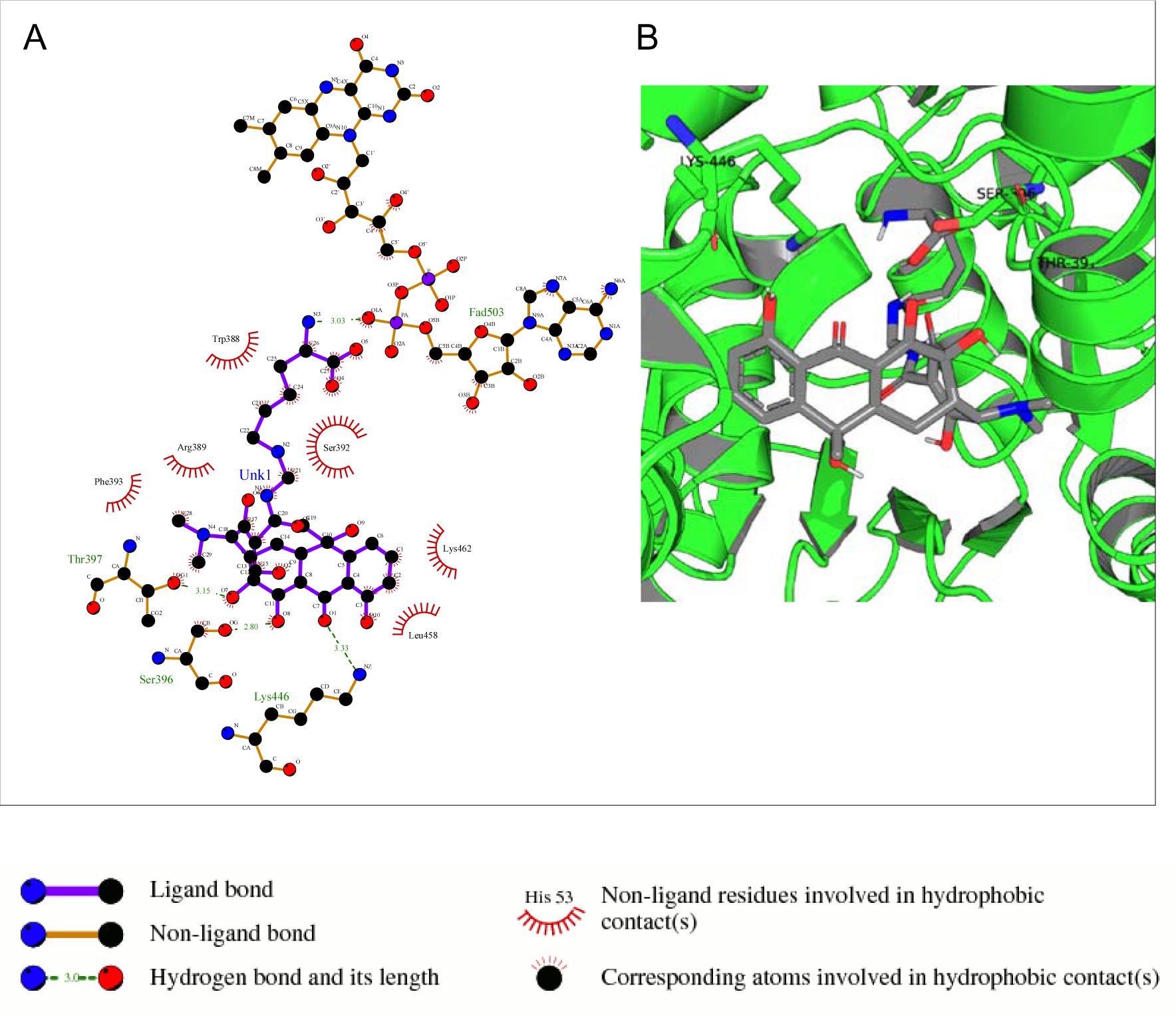
In the above complex, receptor residues Thr-397, Ser-396 (TM12) and Lys-446 (TM17) form hydrogen bonds with the ligand Drugbank138

**Fig.15:**
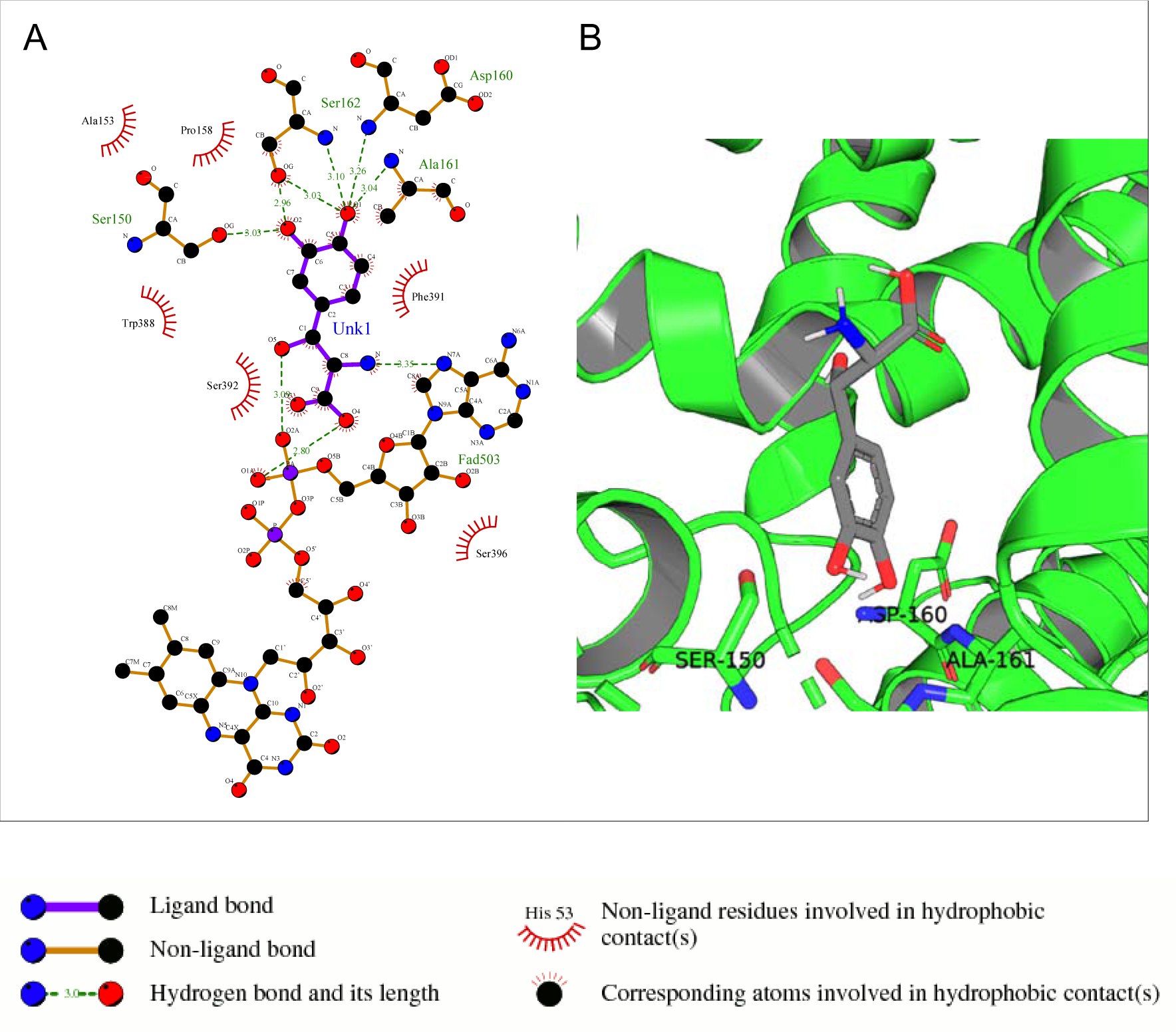
In the above complex (A), residues Asp-160, Ser-162,150 and Ala-161 all within the NADP binding domain form hydrogen bonds with the ligand DrugBank1423.

**Fig.16:**
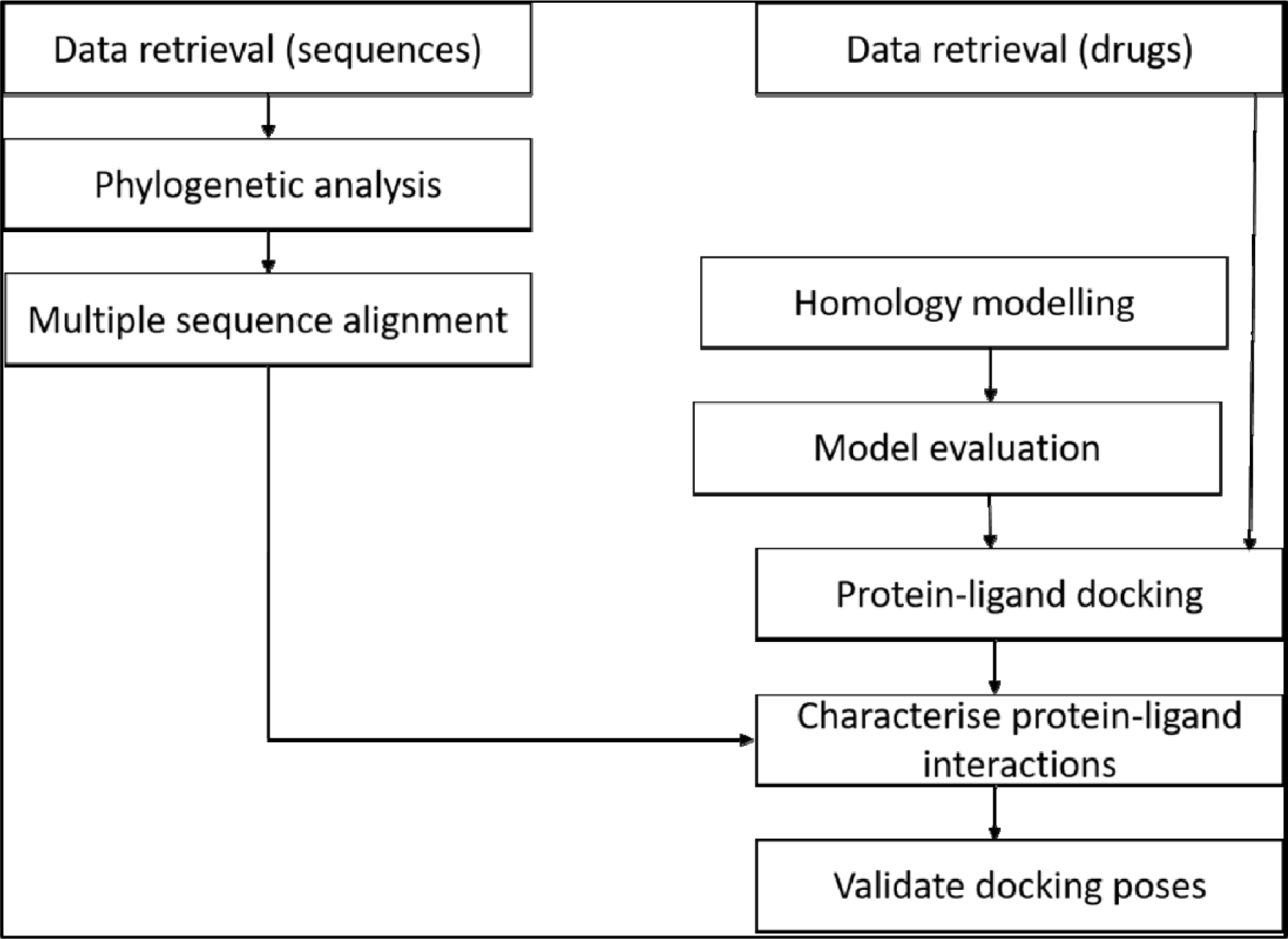
A study design of the methodology. The relationship of data from multiple sequence alignment of related species to human STEAP2 gave insight on key protein residues important in function of the protein to look out for in ligand interaction with the protein during protein-ligand docking.

Ser-372 with the hydrogen bond lies in the 6^th^ TM domain which is part of the iron binding domain and this is shown to be highly conserved from the MSA of only STEAP2 containing species (Fig. 7). It together with many other shown protein amino acids bind with Triptorelin a drug approved for treatment of advanced prostate cancer.

The interacting protein amino acids lie within the TM domains that are highly conserved as shown in the MSA (Fig. 7). These are associated with the iron binding domain and bind to the ligand Leuprolide which is also approved for treatment of advanced prostate cancer.

Some of the listed residues such as Thr-210 are conserved under the NADP domain, the 5^th^ TM domain having residues Glu-390 and Lys-303. The residues together with other form bonds with ligands Drugbank2328 which is 1,2 icosapentoyl-sn-glycero-3-phosphoserine an approved experimental drug for Corona virus-19 disease treatment.

The listed amino acids from the protein that form hydrogen bonds are all conserved highly in the NADP binding domain and the 12^th^ TM domain which is associated with iron binding. The ligand Drugbank2154 is Etelcalcetide a calcimimetic drug used for treatment of hyperparathyroidism.

The mentioned residues in TM12 are associated with the iron binding domain of STEAP2 proteins in the MSA. The ligand Drugbank138 is Lymecycline a second generation tetracycline antibacterial used for broad-spectrum treatment of various bacterial infections.

From the images above, the protein ligand complex of ligand 1543 had the least number of hydrogen bonds between the ligand and protein but had the most number of hydrophobic contacts within the receptor. The summation of these bonds was therefore responsible for the high binding energy associated with the complex. The ligand Drugbank 1423 is Droxidopa a drug used in treatment of Parkinsonism.

Table 3 shows the protein-ligand complexes that met the thresholds of the binding energy.

**Table 3:**
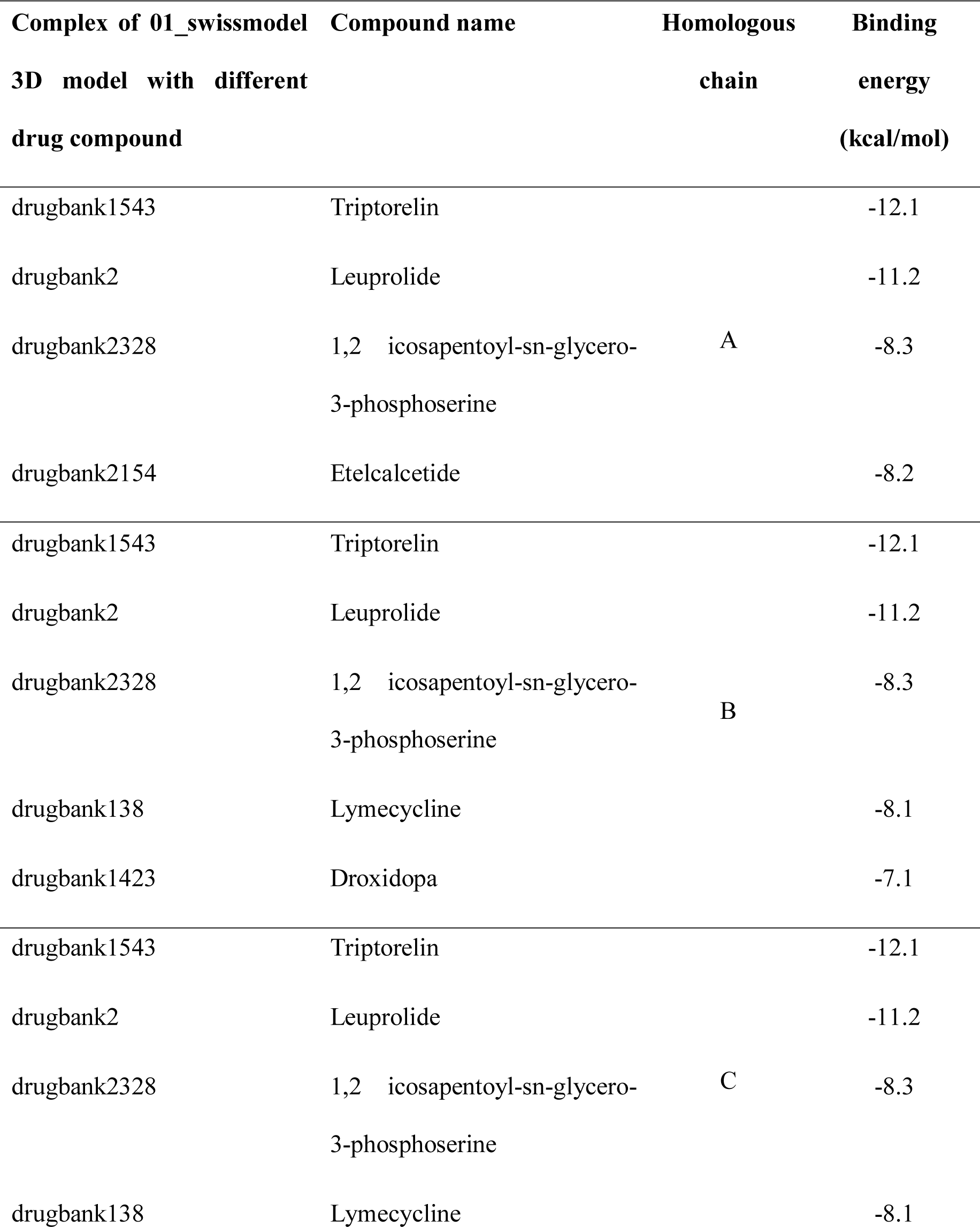
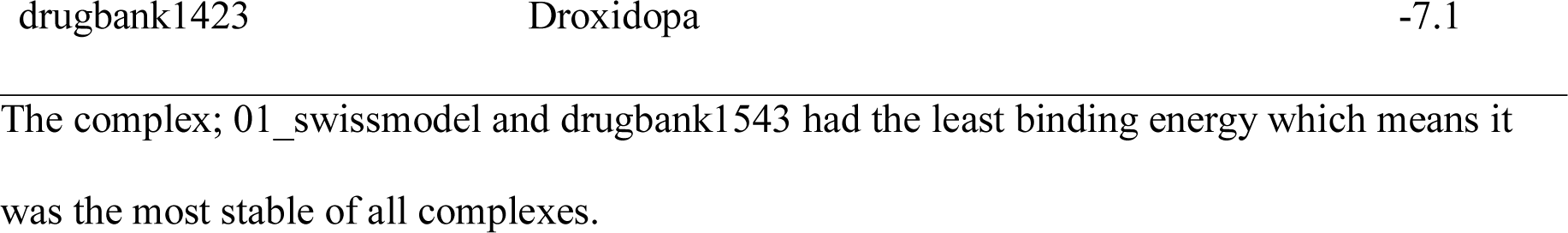
Protein-ligand complexes that met the binding energy threshold with respect to the different chains of the protein.

| **Complex of 01_swissmodel**<br>**3D model with different**<br>**drug compound** | **Compound name** | **Homologous**<br>**chain** | **Binding**<br>**energy**<br>**(kcal/mol)** |
| --- | --- | --- | --- |
| drugbank1543 | Triptorelin | A | -12.1 |
| drugbank2 | Leuprolide |  | -11.2 |
| drugbank2328 | 1,2 icosapentoyl-sn-glycero-3-phosphoserine |  | -8.3 |
| drugbank2154 | Etelcalcetide |  | -8.2 |
| drugbank1543 | Triptorelin | B | -12.1 |
| drugbank2 | Leuprolide |  | -11.2 |
| drugbank2328 | 1,2 icosapentoyl-sn-glycero-3-phosphoserine |  | -8.3 |
| drugbank138 | Lymecycline |  | -8.1 |
| drugbank1423 | Droxidopa |  | -7.1 |
| drugbank1543 | Triptorelin | C | -12.1 |
| drugbank2 | Leuprolide |  | -11.2 |
| drugbank2328 | 1,2 icosapentoyl-sn-glycero-3-phosphoserine |  | -8.3 |
| drugbank138 | Lymecycline |  | -8.1 |

### Docking Validation using independent docking engines

The top 6 mentioned ligands’ binding energy order was re-evaluated using an independent docking engine ; Patchdock and it produced the same order in terms of scores based on the geometric shape complementarity of solutions (29). The scores were ranked in the same order as the binding energy measured by autodock vina

Table 4 shows the top 3 solution of each complex ranked in order of their scores by Patchdock. The area stands for approximate interface area of the complex and atomic desolvation energy is measured according to Zhang *et al.* (30).

**Table 4:** Patchdock docking evaluation of docking scores by Autodock Vina

| STEAP2 com-<br>plex with; | Results'<br>number | Score | Area | Atomic<br>Desolvation<br>Energy | Binding energy<br>measured by<br>Autodock vi-<br>na(Kcal/mol) | Rank |
| --- | --- | --- | --- | --- | --- | --- |
| Drugbank1543 | 1 | 11346 | 1452.70 | -272.92 | -12.1 | 1 |
|  | 2 | 11170 | 1468.30 | -254.68 |  |  |
|  | 3 | 1114 | 1512.50 | -262.71 |  |  |
| Drugbank2 | 1 | 11274 | 1460.40 | -454.04 | -11.2 | 2 |
|  | 2 | 11088 | 1490.30 | -502.35 |  |  |
|  | 3 | 11086 | 1469.60 | -501 |  |  |
| Drigbank2328 | 1 | 11224 | 1375.80 | -421.80 | -8.3 | 3 |
|  | 2 | 10568 | 1383.10 | -501.26 |  |  |
|  | 3 | 10506 | 1321.70 | -462.1 |  |  |
| Drugbank2154 | 1 | 9240 | 1222.30 | -769.57 | -8.2 | 4 |
|  | 2 | 9096 | 1187.60 | -734730 |  |  |
|  | 3 | 9094 | 1132.30 | -453.96 |  |  |
| Drugbank138 | 1 | 7074 | 814.80 | -421.80 | -8.1 | 5 |
|  | 2 | 7016 | 797.00 | -382.80 |  |  |
|  | 3 | 6970 | 841.00 | -373.47 |  |  |
| Drugbank1423 | 1 | 3300 | 358.90 | -133.28 | -7.1 | 6 |
|  | 2 | 3294 | 357.70 | -125.81 |  |  |
|  | 3 | 3294 | 363.30 | -122.28 |  |  |

The exact docking poses from autodock vina could not be reproduced by patchdock despite maintaining the same ranking order. Another docking engine; Autodock tools could not reproduce the exact docking poses.

## Discussion

The development of a more specific drug for the treatment of prostate cancer not only reduces side effects of the current regimen but increases potency and effectiveness of the candidate. With a modelled homologue and docking process of a pool of approved candidates from the drug bank, a number of drug compounds that formed significant interactions with the homology model gave insights of some of the current drugs that could be repurposed for the treatment of prostate cancer.

The best homology model of STEAP2 was generated by SwissModel as shown by the results from assessment by the 5 evaluation tools used (Fig.8-A). SwissModel, in a study by Wallnar has been shown to accurately model homologues whose templates are of as low as 40% similarity (31). This suggests that the modelling engine is reliable.

On model analysis, a lower score below 100% in the favoured region on a Ramachandran plot shows that there are residues whose phi and psi angles are not sterically possible when modelled (32). A lower percentage of residues in the favoured region therefore indicates a poor homology model as shown for models from Robetta and I-Tasser.

From the docking results, ligand Drugbank1543 had the least binding energy and highest affinity from a selected set of 2466 ligands. Having the least binding energy means the complex is more naturally stable due to affinity between the two constituent molecules with no or less external energy applied. The ligand formed its hydrogen bond with a residue Ser372 in the transmembrane domain of the receptor molecule as most of its other non-hydrogen bond interactions.

From a key of the drugbank, Drugbank1543, accession number DB06825, generic name Triptorelin is a synthetic decapeptide gonadotrophin releasing hormone (GnRH) agonist. The drug is proven for treatment of advanced stage prostate cancer as palliative care to reduce sequelae of advanced disease such as compression of the spinal cord (33). With the results from this study, this drug could potently be used to treat early stage disease targeting STEAP2 that is shown to be responsible for disease progression.

The study by Ravenna *et al.* demonstrates that Triptorelin has an inhibitory-stimulatory effect on Lymph Node Carcinoma of the Prostate (LNCaP) cells at lower and high doses respectively. In the experiment, in Pc3 cell lines, only low affinity but high capacity receptors were expressed as a biological reaction to the drug. Both low and moderately high affinity with high and low capacity receptors respectively were expressed in LNCaP cells. (34). This shows the potency of Triptorelin and its binding energy in the treatment of prostate cancer at different stages. Since STEAP2 is present in both cell types and given the binding relationship between the protein and drug, this study suggests a mode of action of the latter.

Drugbank2 accession number DB00007 generic name Leuprolide is a synthetic 9 residue peptide analogue of GnRH with the same mode of action as Triptorelin and is equally indicated for treatment of advanced prostate cancer (35,36). It has been shown to have a quicker onset of action in achieving serum testosterone levels equivalent to medically castrated men compared to Triptorelin (37). The findings from this study suggest STEAP2 as a target route for action.

Data from the protein atlas places brain tissue (pituitary gland, cerebral cortex and basal ganglia) third after the ovary and vagina. In men where the ovary and vagina are absent, brain tissue becomes second after the prostate though with a significant difference in tissue STEAP2 expression. This could account for some of the potency of the GnRH receptor analogue agonists in treating prostate cancer as mentioned above.

Drugbank2328, accession number DB14096, 1,2 icosapentoyl-sn-glycero-3-phosphoserine is an approved experimental drug for the treatment of Corona Virus Disease-19 (COVID-19) (38).Drugbank138, accession number, DB00256, Lymecycline is a broad spectrum second generation tetracycline antibacterial used for treatment of susceptible bacterial infections such as acnes (39). Drugbank1423, accession number DB06262, generic name, Droxidopa a precursor for noradrenaline and is in the treatment of Parkinsonism (40). Drugbank2154, accession number DB12865, generic name Etelcalcetide is a calcimimetic drug used for the treatment of secondary hyperparathyroidism in patients having haemodialysis. From the findings of this study, these drugs could treat prostate cancer by binding to and inhibiting the catalytic role of STEAP2 that is responsible for disease progression. Further investigation on their role in treatment of prostate cancer is still warranted.

The exact docking poses could not be reproduced by autodock tools probably because the engine requires use of a single chain of the homo-3-mer for the protein ligand docking vis-à-vis the 3 identical chains used in the docking by autodock vina. The coherence in ranking order of protein ligand complex stability between autodock vina and patchdock, however, suggests the complexes are rightfully ordered.

Due to time and cost limitations we were unable to successfully validate the docking poses of the top ligands using different and independent docking engines. We recommend that future work should include reproducing the docking poses using other commercial and/or free docking engines in addition to molecular docking simulations. The docking engine used (Autodock vina) is, however, documented to be one of the best at the time as shown in the study by Sousa *et al*. (41). Additionally, we also recommend further analysis in terms of *in-vitro* assays to validate the results from this *in-silico* study on growth and viability changes in the presence of these drugs.

## Materials and methods

### Study design

The study involved retrieval of STEAP2 related protein sequences from the protein data bank from a protein BLAST search. The selected sequences were used for an MSA to phylogenetic analysis to determine the most related species to human STEAP2 protein. An MSA was used to determine highly conserved residues important in function of STEAP2 protein.

For molecular docking, a set of FDA approved drugs were retrieved from the drug bank and these were used as ligands for protein-ligand docking. The protein (STEAP2) 3 Dimension structure was acquired from homology modelling since it did not exist as an approved structure in the protein data bank. The homology model, after being evaluated was used for molecular docking with the drugs from the drug bank and the results of each were ranked.

The Protein-ligand interactions were characterised using ligpots and protein interacting residues were correlated with the those earmarked for protein function from the MSA and evaluated using independent docking engines. This entire methodology is summarised in Fig.16.

### STEAP2 relative tissue expression

To determine the expression of STEAP2 protein in different tissues so as to ascertain its specificity as a drug target, an expression analysis was done using the Human Protein atlas (https://www.proteinatlas.org/ENSG00000157214-STEAP2) that showed its relative expression in over 50 different body tissue in addition to the prostate (18).

### Data retrieval

The amino acid sequence of the query protein *H. sapiens* STEAP2 (Uniprot accession number: Q8NFT2) was obtained in FASTA format from UniProt (42). This sequence was used in the protein BLAST search for homologous sequences in the PDB.

### STEAP2 Multiple Sequence Alignment and phylogenetic analysis

The MSA was done to determine the essential residues for the metalloreductase activity of STEAP proteins. A protein BLAST (BLASTp) search of the reference proteins in the PDB was done with the algorithm parameter of maximum target sequences to return adjusted to 5000. 30 sequences of the 5000 returned were selected for MSA. These were selected basing on percentage similarity in bins of 0-40, 41-60, 61-80 and 81S-100%. A number of sequences, was selected from each range to have a uniform selection and also minimise bias in the phylogenetic analysis. (Table 5).

**Table 5:**
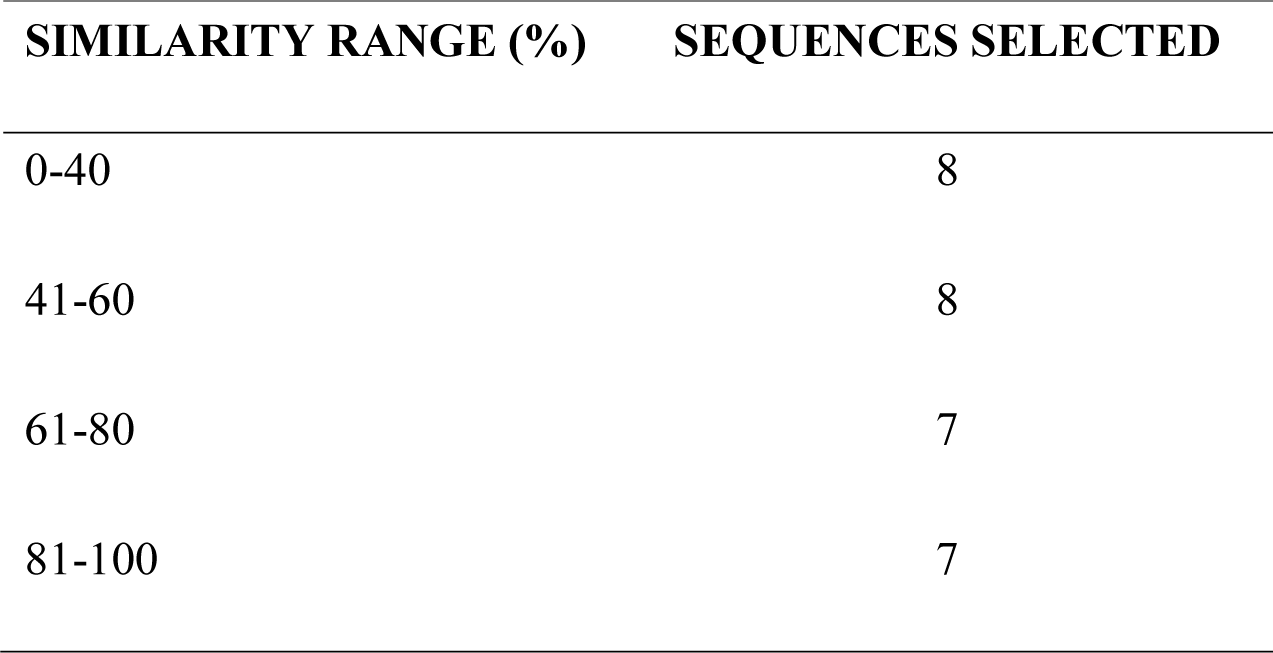
Sequences selected for phylogenetic analysis of STEAP2 by MSA.

From the table, the 0-40 and 41-60 similarity ranges took an extra hit so as to ensure that more distant hits from the query sequence were considered for confidence purposes.

All selected sequence files were downloaded in FASTA format and used in an MSA utilizing Muscle in MEGAX so as to determine the conserved regions between the different metalloreductases from 30 different species with different similarities to STEAP2 (21).

In MEGAX, a new protein alignment was created. In phylogeny, the bootstrap method was used, an LG+G+I model selected and the number of bootstraps set to 1000. The rest of the settings were left as default. A dendogram showing a phylogenetic relationship of STEAP2 was generated (Fig. 6 and 7).

### Homology modelling

Using data from MSA, we were able to identify possible similar sequences from the 80-100% similarity range to the STEAP2 sequence giving direction for hits that might already have a determined 3D structures and would act as templates for homology modelling.

### Web-based homology modelling engines

*H. sapiens* STEAP2 (Uniprot accession number: Q8NFT2) FASTA sequence was used as a query in the different modelling engines that were best ranked by the Continuous Automated Model Evaluation (CAMEO) website and one other from the Critical Assessment of Protein Structure Prediction (CASP) website (I-TASSER) to search for similar sequences for homology modelling (43,44). The different homology modelling engines BLAST searched the PDB for similar sequences that have approved 3D structures. The similarity of retuned results from the BLAST was shown in terms of coverage of the query sequence, the percentage identity and the E-value. Also, the organism that was the source of the sequence and the PDB identification of each result was shown. The best hit’s resolution and method of structural determination were obtained from the protein data bank(45).

The best sequence in all but those of RaptorX and I-TASSER were manually chosen for homology modelling. I-TASSER automatically chose the best nine sequences and made homology models of STEAP2 using each and the best ranked result (homology model 1) was chosen as the final homology model. RaptorX automatically chose its best template and the showed its homology model as a result at the end of the process.

The top templates by the rest of the engines were used to model STEAP2 using Modeller and ProMod3 (46,47). The homology models were downloaded as PDB files and visualised using PyMol and Chimera (48,49)

### Model Evaluation of homology models

The homology models from the different modelling engines were downloaded in PDB format and evaluated using different structural model evaluation tools i.e., ProSa, QMEAN, Rampage, DOPE scores and RMSD. Ramachandran Plot Server (URL: https://zlab.umassmed.edu/bu/rama/index.pl was used to generate Ramachandran plots of the homology models. Using PyMol, the stoichiometry of the homology model was determined by chain coloration and critically looking at structures to identify any non-conformities such as long loops or knots (50) RMSD with the 6HCY (human STEAP4) template was calculated. by use of PyMol alignment tool to align the template with the homology models to determine average structural deviations of the homology model from that of the used template.

The DOPE score of each homology model was calculated by Modeller. Z-DOPE scores were calculated for individual homology models using Modeller which was installed using Conda from commands from the site https://anaconda.org/salilab/modeller and it was used to calculate the normalised z-DOPE score of each homology model(47,51)

### Molecular Docking

With the best model selected from 6 STEAP2 homology models after evaluation, the homology model by Swissmodel was used for flexible ligand, rigid receptor protein-ligand docking using a group of approved drug compounds downloaded from the drug bank. These compounds were downloaded as a SANCDB file of 2466.

The docking was done using AutoDock Vina. The ligand and receptor molecules were prepared using prepare_ligand4.py and prepare_receptor4.py respectively by command line on Linux terminal. The search box in the configuration file was set to dimensions x,y,z (105 by 105 by 105 Å, with a centre at: x=122.224Å, y=118.600Å, and z=102.658Å).

The results of molecular docking were characterised by 3D proximity to a central residue termed as “Euclidian distance”-which is the length of a line between two points in geometrical space which can be 2 or 3D and binding energy of the ligand to the receptor molecule. All results were screened and a binding energy threshold of less than or equal to -7 and Euclidian distance of 50Å were set. The complexes that met the selection criteria were then considered as the candidates for further analysis.

The entire protein was covered by the search box during molecular docking; a method also known as blind docking such that the docking is not biased to any region of the receptor.

### Characterising Protein-Ligand Interactions

To determine the nature of the bonds formed between STEAP2 and the different ligands, Ligplot which is a program used to generate 2D representations of the nature of bonds formed between protein receptor residues and ligand atoms was used for this task (52). The program showed the hydrogen bond interactions in green dotted lines with their respective lengths in angstroms and the different interacting protein residues in an arc shape with spokes radiating towards the atoms they are in contact with. The interactions not only accounted for hydrogen bonds but all hydrophobic interactions between the two molecules (52).

### Docking validation using independent docking engines

The order of binding energy was validated using patchdock server and autodock tools. (29,53). The receptor pdb file and ligand pdbqt files were uploaded on pactchdock server and results were downloaded via a link from the webserver. In autodock tools, the protocal in the tutorial by Huey *et al.* was followed (53)

## Supporting information

Table containing details on sequences' details used for MSA

Fasta file for STEAP2

## Acknowledgments.

Special appreciation to the Department of Biomolecular Resources and Biolaboratory sciences for the guidance and resources used in executing this work.

